# Local brassinosteroid biosynthesis enables optimal root growth

**DOI:** 10.1101/2020.11.23.391474

**Authors:** Nemanja Vukašinović, Yaowei Wang, Isabelle Vanhoutte, Matyáš Fendrych, Boyu Guo, Miroslav Kvasnica, Petra Jiroutová, Jana Oklestkova, Miroslav Strnad, Eugenia Russinova

## Abstract

Brassinosteroid hormones are indispensable for root growth and they control both cell division and cell elongation through the establishment of an increasing signaling gradient along the longitudinal root axis. Because of their limited mobility, the importance of brassinosteroid distribution for achieving the signaling maximum is largely overlooked. Expression pattern analysis of all known brassinosteroid biosynthetic enzymes revealed that not all cells in the *Arabidopsis thaliana* root possess full biosynthetic machinery and completion of biosynthesis relies on cell-to-cell movement of the hormone precursors. We demonstrate that brassinosteroid biosynthesis is largely restricted to the root elongation zone where it overlaps with brassinosteroid signaling maxima. Moreover, optimal root growth requires hormone concentrations, low in the meristem and high in the root elongation zone attributable to an increased biosynthesis. Our finding that spatiotemporal regulation of hormone synthesis results in a local hormone accumulation provides a paradigm for hormone-driven organ growth in the absence of long-distance hormone transport in plants.

---

Brassinosteroids (BRs) are a group of steroidal phytohormones essential for plant growth and development^1^. Mutants deficient or insensitive to BRs exhibit a range of growth defects, including dwarfism^2,3^, photomorphogenesis in the dark^4^, altered stomatal development^5,6^ and reduced male fertility^7^. Because of their importance for plant development, BRs have attracted much research attention in the past two decades and therefore, BR signaling is one of the best-characterized signal transduction pathways in *Arabidopsis thaliana*^8^. The BR signaling cascade starts with a direct binding of a BR ligand to the extracellular domain of the plasma membrane (PM)-localized receptor kinase BRASSINOSTEROID INSENSITIVE1 (BRI1) or its close homologs BRI1-LIKE1 (BRL1) and BRL3^9,10^. Subsequently, transphosphorylation events between the kinase domains of BRI1 and its coreceptors of the SOMATIC EMBRYOGENESIS RECEPTOR KINASE (SERK) family occur and fully activate the receptor complex^11^. Further phosphorylation-dephosphorylation steps lead to dephosphorylation of the key transcription factors BRASSINAZOLE RESISTANT1 (BZR1) and BR INSENSITIVE EMS SUPPRESSOR1 (BES1)/BZR2 and their translocation into the nucleus^12^, where they stimulate or repress thousands of specific genes required for optimal BR responses^13^.

BRs are derived from campesterol that is converted to brassinolide (BL), the most potent and final product of the BR biosynthetic pathway, through a series of reactions, including reduction, hydroxylation, epimerization and oxidation^1^. Extensive research by means of feeding experiments with labeled intermediates^14^ have established a biosynthetic grid with multiple possibilities through which BL could be synthetized. Nevertheless, evidence on the enzymatic substrate specificity and intermediate content analysis of the mutants^15,16^ made it possible to propose a succession of biosynthetic steps and positioning of the enzymes in a somewhat linear order (Supplementary Fig. 1a)^17^. With the exception of DE-ETIOLATED2 (DET2), which is a steroid 5α-reductase^18^, all known enzymes of the BR biosynthetic pathway belong to the family of the cytochrome P450 (CYP450) monooxygenases^1^.

Hormone levels at any given site are determined by the relative rates of *de novo* synthesis and catabolism, conjugate synthesis, and hydrolysis, and, finally, transport from the site of synthesis to the site of action^19^. Studies on BR transport in different plant species have demonstrated the absence of hormonal organ-to-organ mobility^20^, thus reinforcing the importance of the regulation of the biosynthesis and catabolism rates through transcriptional feedback loops^1^. Increased transcript levels of BR biosynthetic genes are usually correlated with young developing tissues and higher BR content^21–23^. Nonetheless, a high-resolution spatiotemporal map of BR biosynthesis is still lacking.

The *Arabidopsis* root has been used extensively as a model organ for BR signaling^24^ and plant organ growth studies^25^. The apically positioned meristem is a region of actively dividing cells that gradually, through longitudinal expansion, leave the meristem region and enter the division-free elongation zone where they reach their mature size. The meristem size and, thus, the number of growing cells are critical parameters for optimal root growth^26^. As BRs regulate cell division^27^ and control cell elongation^28^, they have a paramount importance for root meristem size determination. Elevated BR signaling results in premature elongation and differentiation of meristematic cells and has a negative effect on meristem size and root growth^27^, possibly due to the existing BR signaling gradient along the longitudinal axis of root tips, with a peak in the elongation zone^29^. However, the factors that determine the BR signaling gradient in the *Arabidopsis* root remain elusive.

Here, through localization studies, we show that not every root cell type expresses all BR biosynthetic enzymes and a completion of BR biosynthetic pathway relies on cell-to-cell movement of the hormone precursors. Moreover, most of BR biosynthetic enzymes reach their expression maxima in the root elongation zone to ensure high hormone levels required for the BR signaling maintenance. Thus, local BR biosynthesis plays a crucial role in the formation of the signaling gradient along the longitudinal root axis, enabling the optimal meristem activity and timely cell elongation.

## Results

### BR Biosynthetic Enzymes Are Not Expressed in Every Cell Type of the *Arabidopsis* Root

*Arabidopsis* mutants defective in BR biosynthesis and signaling display severe phenotypic defects which include compact rosette with dark-green curled leaves and short roots with an increased diameter (Supplementary Fig. 1b-f). To gain insight into the importance of the BR synthesis and distribution for growth and morphology of *Arabidopsis* roots, we carried out localization studies for all known enzymes in the pathway (Fig. 1a). Genomic fragments of each BR biosynthetic gene, including at least 2,000-bp promoter regions upstream from the start codon, were fused with the coding sequence of the green fluorescent protein (GFP) and introduced into the corresponding mutant backgrounds. All tested protein fusions were functional, because they complemented the mutant phenotypes and their expression patterns were identical in multiple independent transgenic lines (Supplementary Fig. 2a,b). First, we examined the enzyme expression patterns in the *Arabidopsis* root meristem. The expression of CPD-GFP, DET2-GFP, BR6OX1-GFP and GFP-BR6OX2 was confined to the inner tissues of the root apical meristem (Fig. 1a). The CPD-GFP expression occurred in procambial and central columella cells (Fig. 1a,c), whereas BR6OX1-GFP and GFP-BR6OX2 were expressed mainly in the endodermis and pericycle (Fig. 1a,g,h). A weak GFP-BR6OX2 signal was observed in young cortical cells (Fig. 1i). The expression of DWF4-GFP and DET2-GFP in the apical meristem was somewhat broader and included additional cell layers for DET2-GFP (Fig. 1a,d) and a few cell types for DWF4-GFP, with an expression maximum in the epidermis (Fig. 1a,b). The expression domain of ROT3 and CYP90D1 was more widespread and incorporated procambial, cortical and columella cells (Fig. 1a,e,f). Gene expression reporter lines in which the promoters of the BR biosynthetic genes were fused to N-terminal nuclear localization sequence (NLS)-GFP showed largely similar expression patterns (Supplementary Fig. 2c,d).

**Fig. 1.**
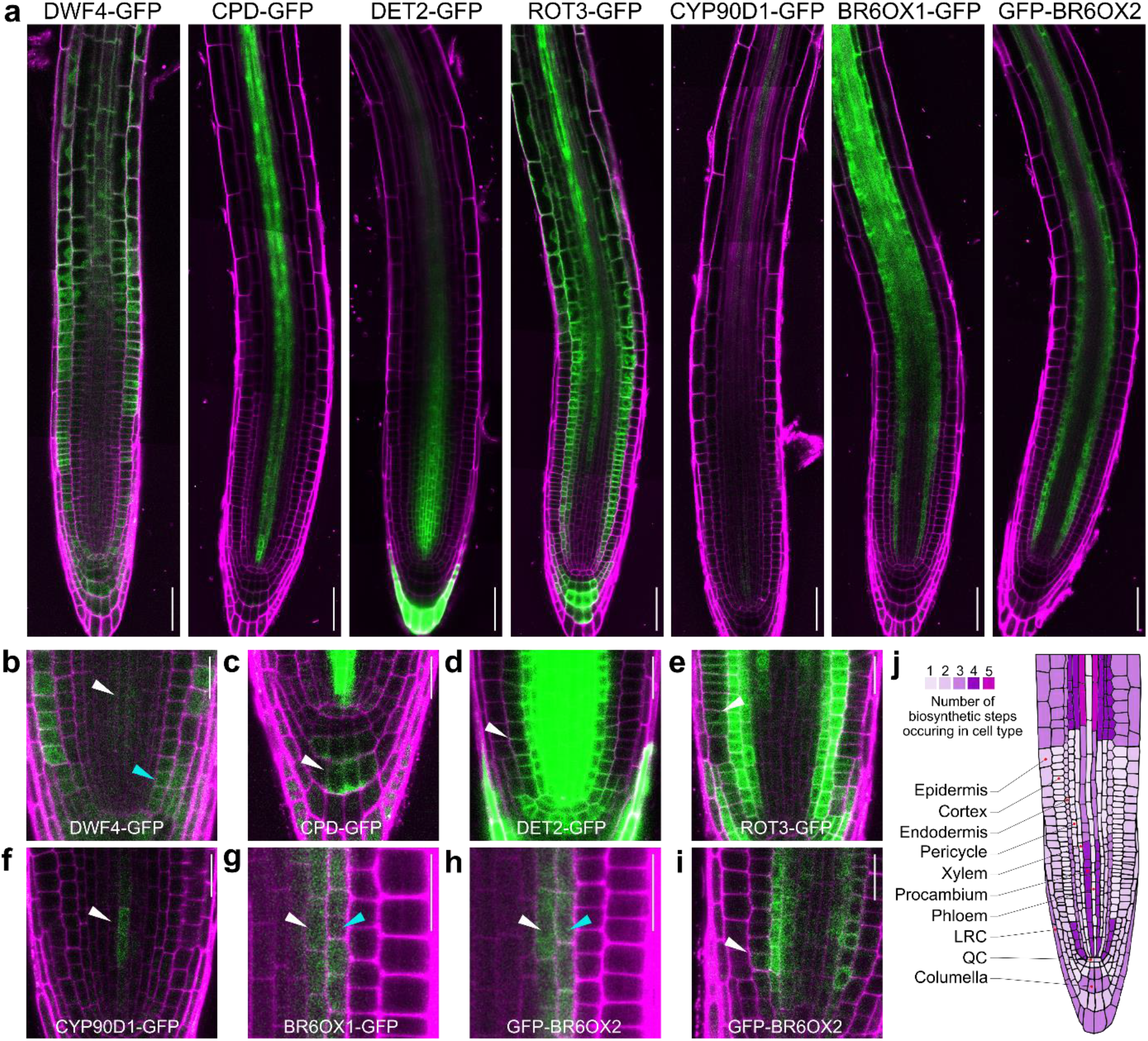
Expression patterns of BR biosynthetic enzymes in the *Arabidopsis* root. **a,** Roots tips of 6-day-old seedlings expressing DWF4-GFP, CPD-GFP, DET2-GFP, ROT3-GFP, CYP90D1-GFP, BR6OX1-GFP and GFP-BR6OX2 under the control of their native promoters. **b,** Weak DWF4-GFP signal in procambial (white arrowhead) and cortical cells (blue arrowhead). **c,** Limited expression of CPD-GFP in the central columella region (white arrowhead). **d,** Extended DET2-GFP expression at the very root tip of the root, including young cortical cells (white arrowhead). **e,** Occurrence of ROT3-GFP signal in the epidermis (white arrowhead) in addition to the dominant expression in the cortex. **f,** Weak CYP90D1-GFP expression in procambial cells in the root tip near the quiescent center (white arrowhead). (**g** and **h**) BR6OX1-GFP (**g**) and GFP-BR6OX2 (**h**) expression limited to the pericycle (white arrowheads) and endodermis (blue arrowheads). (**i**) GFP-BR6OX2 expression in young cortical cells. For panels (B-H) GFP signal brightness and contrast were adjusted to make it more visible in cell types of interest. (**j**) Schematic representation of the *Arabidopsis* root tip with color code displaying the number of BR biosynthetic steps occurring in a certain cell type. Only procambial cells in the elongation zone of the root express all BR biosynthetic enzymes. LRC, lateral root cap; QC, quiescent center. Scale bars represent 50 μm in (**a**) and 20 μm in (**b-i**). Roots were stained with propidium iodide.

Our localization studies indicate that most cells in the *Arabidopsis* root meristem do not possess all enzymes needed to complete the BR biosynthetic pathway (Fig. 1j), implying that intermediates have to be exchanged between the cells in order to finalize hormone synthesis.

### BRs Can Move Over Short Distances in the *Arabidopsis* Root

Although BRs are not transported over long distances^20^, the spatially restricted expression of BR biosynthetic enzymes in the *Arabidopsis* root (Fig. 1a), in contrast to the broad expression domain of the BRI1 receptor^30^, implies an obligatory BR movement. To test this hypothesis, we used the *CPD* gene, which has the most restricted expression domain in the root meristem (procambial cells) of all BR biosynthetic genes, and expressed it ectopically in the endodermal cell files by means of the endodermis-specific *SCARECROW* (*SCR*) promoter^31^ (Fig. 2a). Ectopic expression of CPD-mCHERRY in the endodermis complemented the root growth defects of the *cpd* mutant (Fig. 2b,c) and restored the normal root meristem architecture (Fig. 2d,e and Supplementary Fig. 3a,b). Restoration of the BR signaling in endodermal cell files had previously been shown to be insufficient to rescue the root growth defects of the *bri1* signaling mutant^29,32^. As the *bri1* and *cpd* mutants have similar phenotypes due to the lack of BR signaling and BR hormones, respectively, our findings demonstrate that the restoration of the BR biosynthesis, but not of its signaling, in endodermal cell files can complement the *cpd* mutant phenotype, probably because of the mobile nature of the hormone or its intermediates.

**Fig. 2.**
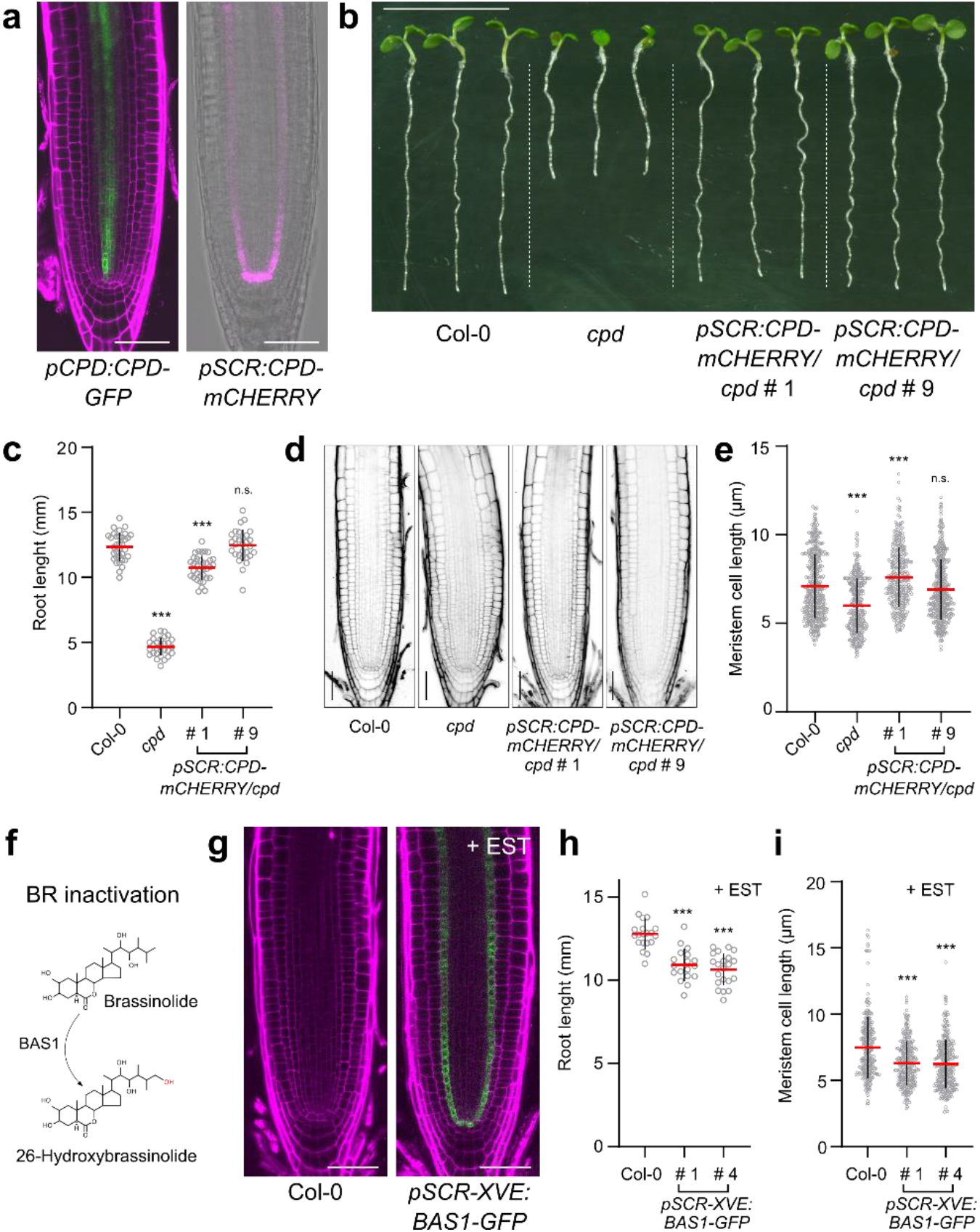
BRs can move over short distances in the *Arabidopsis* roots. **a,** CPD-GFP expression restricted to procambial cells of the *Arabidopsis* root when expressed under its native promoter (left panel). Ectopic expression of CPD-mCHERRY from the endodermis-specific SCR promoter in the *cpd* mutant (right panel). **b,** Six-day-old *cpd* mutant seedlings expressing *pSCR:CPD-mCEHRRY*. Two independent lines are shown. **c,** Quantification of the root lengths of the lines shown in (**b**). Six-day-old plants were transferred to fresh medium and root growth was measured after 3 days, *n* ⩾ 29 roots analyzed. **d,** Root meristem architecture of *Arabidopsis* wild type (Col-0), *cpd* and transgenic lines expressing *pSCR:CPD-mCHERRY* in the *cpd* mutant. **e,** Quantification of cortical meristem cell length (first 20 cells of individual roots) of lines expressing *pSCR:CPD-mCHERRY* in the *cpd* mutant, *n* ⩾ 17 roots analyzed for each genotype. **f**, BAS1 catalyzing the conversion of BRs to inactive C-26 hydroxylated forms. **g,** Root tips of 7-day-old Col-0 and *pSCR-XVE:BAS1-GFP* seedlings grown in the presence of 10 μM β-estradiol (EST) for 3 days. BAS1 expression is limited to the endodermal cell layer. **h,** Quantification of the root lengths of two independent lines expressing *pSCR-XVE:BAS1-GFP*. Four-day-old plants were transferred to fresh medium containing 10 μM β-estradiol and the primary root length was measured after 3 days, *n* ⩾ 18 roots analyzed. **i,** Quantification of cortical meristem cell length (first 20 cells of individual roots) of lines expressing *pSCR:CPD-mCHERRY* in the *cpd* mutant, *n* ⩾ 18 roots analyzed for each genotype. Grey circles, red wide bars and black whiskers in the plots represent individual values, means and SD, respectively. The significant differences between transgenic lines and the Col-0 control were determined by one-way analysis of variance (ANOVA) and Dunnett’s multiple comparisons tests. *** *P* < 0.001, ** *P* < 0.01 and * *P* < 0.05). Roots were stained with propidium iodide. Scale bars represent 50 μm in (**a**, **d** and **g**) and 1 cm (**b**)

To further test our hormone mobility hypothesis, we blocked the BR movement in the root by employing the cytochrome P450 monooxygenase PHYB-4 ACTIVATION-TAGGED SUPPRESSOR1 (BAS1) that catalyzes C-26 hydroxylation of BRs and inactivates them (Fig. 2f)^33^. BAS1 is expressed in the outer tissues of the root (lateral root cap and epidermis)^29^ and its overexpression causes typical BR-related phenotypes (Supplementary Fig. 3c,d)^34^. To inactivate BRs that are either synthetized or pass through this centrally positioned tissue, we ectopically expressed BAS1-GFP in the endodermis in an inducible manner (Fig. 2g). Estradiol-inducible expression of BAS1-GFP under the *SCR* promoter hampered root growth and altered the root meristem architecture in transgenic plants, in a manner typical for BR deficient mutants (Fig. 2h,i and Supplementary Fig. 3e-g). These findings demonstrate that BR intermediates and, possibly, the bioactive BRs, brassinolide and castasterone, can be exchanged between neighboring root cells.

### BR biosynthetic Genes Expression and BR Signaling Maxima Overlap in the Root Elongation Zone

While examining the localization of the BR biosynthetic enzymes, we noticed that the DWF4 expression expanded in the elongation zone and included all cell types (Fig. 3a, Supplementary Fig. 4a and Supplementary Video 1). Similarly, the BR6OX1-GFP expression, although still very prominent in the inner tissues, increased and included epidermal and cortical cells of the elongation zone (Supplementary Fig. 4a). Finally, the ROT3-GFP expression that was mostly confined to the cortical cells in the apical meristem also extended to the epidermal cells and peaked in the stele of the elongation zone (Fig. 3A, Supplementary Fig. 4a and Supplementary Video 2). Besides the expansion of the expression domains, the majority of the transgenic lines, except for DET2-GFP and GFP-BR6OX2, showed a clear increase in expression levels along the longitudinal root axis, with maxima in the elongation zone. This was very prominent in the case of CPD-GFP (Fig. 3A, Supplementary Fig. 4b and Supplementary Video 3). Surprisingly, when BR6OX2 was tagged with GFP at the C-terminus and expressed under the transcriptional control of its native promoter, it also showed increase of expression in the root elongation zone (Supplementary Fig. 4c). These findings are consistent with publicly available expression datasets^35^ (Supplementary Fig. 4d) and together hint at an enhanced hormone production in the root elongation zone.

**Fig. 3.**
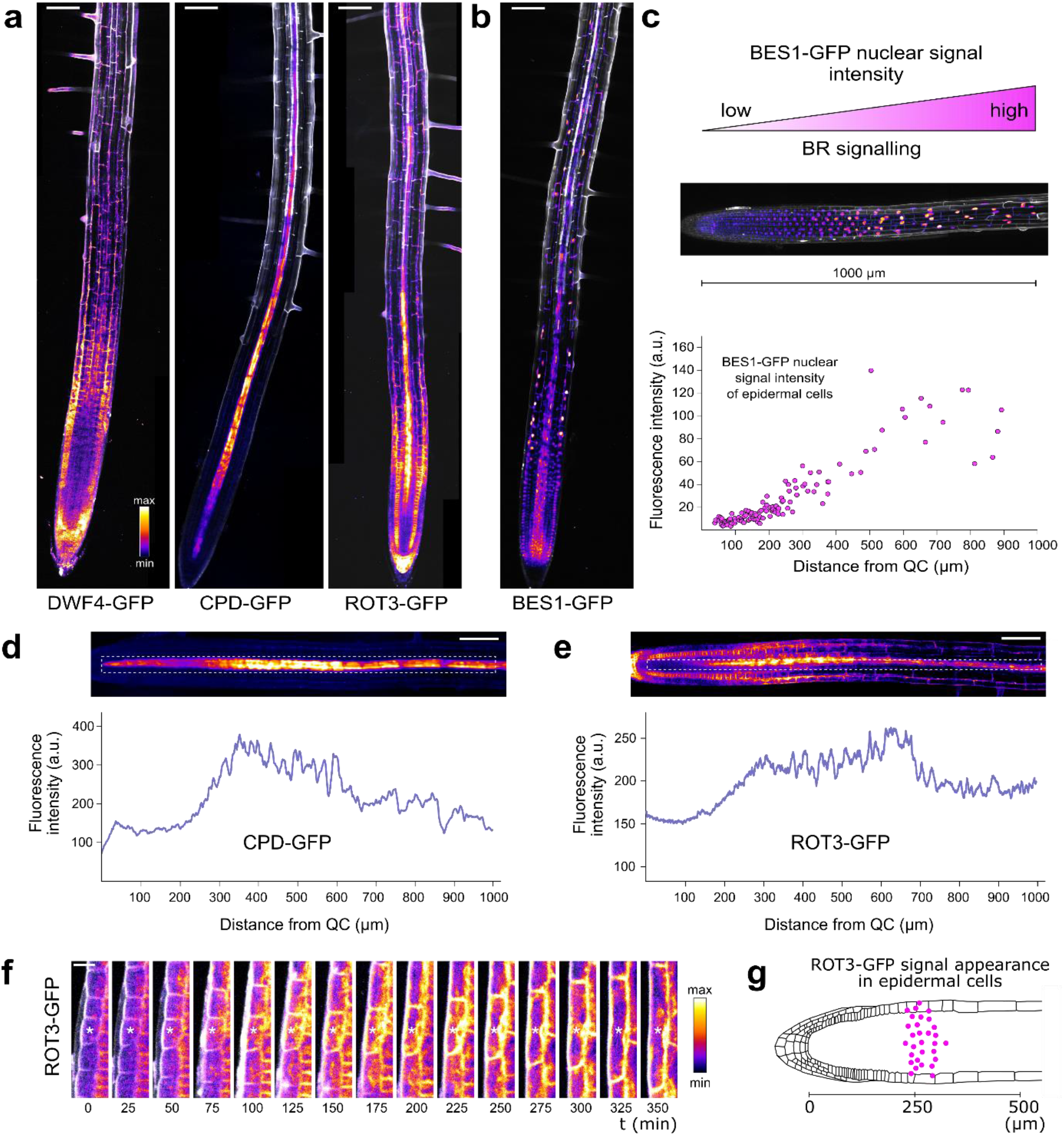
BR signaling and synthesis maxima overlap in the *Arabidopsis* root. **a,** Six-day-old *Arabidopsis* roots expressing DWF4-GFP, CPD-GFP and ROT3-GFP under the control of their native promoters. **b,** Six-day-old *Arabidopsis* root expressing BES1-GFP under the control of its native promoter. **c,** Increase in the BES1-GFP nuclear fluorescence intensity used as readout of active BR signaling. The scatter diagram shows the nuclear fluorescence intensities of BES1-GFP in epidermal cells of a single root in relation to their distance from the quiescent center (QC). **d, e,** Fluorescence intensity profiles of CPD-GFP (**d**) and ROT3-GFP (**e**) along the root elongation zone. The GFP signals were measured along the delimited dashed frames and plotted in the graphs below. The average values of three roots for each line are shown. **f,** Time series analysis of a single epidermal cell expressing ROT3-GFP (white asterisk). The ROT3-GFP signal appears in the transition zone and reaches its peak as the cell enters the elongation zone. **g,** Profile of the ROT3-GFP signal appearance in epidermal cells of the root. Ten time points, 25 min apart, of three independent roots are shown. Scale bars represent 100 μm in (**a**, **b**, **d** and **e**) and 20 μm in (**f**).

The nuclear accumulation of the fluorescently tagged BES1 and BZR1 transcription factors had been used as a readout for active BR signaling^29^. In accordance with the role of BRs in cell elongation, the BR signaling levels show a gradient along the longitudinal root axis with a maximum in the transition-elongation zone^29^, but the regulatory mechanisms establishing this gradient are unclear. We hypothesized that the longitudinal control of the BR biosynthesis is crucial in fueling the BR signaling maxima. To verify this assumption, we compared the BR biosynthetic gene expression patterns with the localization of the BES1-GFP reporter. The nuclear BES1-GFP signal in epidermal and cortical cells started to increase at 200-400 μm from the quiescent center (QC) and reached a peak in the elongation zone (Fig. 3b,c and Supplementary Video 4), as previously reported^29^. Similarly, the fluorescence intensities of both the CPD-GFP and ROT3-GFP signals increased approximately two-fold in the region 200-400 μm from the QC, correlating with the highest BES1-GFP nuclear signal accumulation (Fig. 3d,e). A time-lapse imaging of epidermal cells expressing ROT3-GFP further supported these observations (Fig. 3f). Young epidermal cells showed no or a weak GFP signal while entering the transition zone and as the expansion proceeded, expression started to increase, reaching a peak when the cells entered the elongation zone (Fig. 3g).

Taken together, the expression of BR biosynthetic genes in *Arabidopsis* roots peaks in the elongation zone, as a consequence of an increase in the expression levels and broadening of the expression domain into additional cell files. Notably, this region of the root coincides with the BR signaling maxima domain, implying that the hormone biosynthesis determines the local levels of the signaling responses.

### The Root Elongation Zone Has Higher Demands for BRs than the Root Meristem

Exogenous BL has been shown to restore BR signaling in the *dwf4* mutant root^29^, hence ruling out the importance of local BR biosynthesis for root growth, but the root growth rates and root meristem morphology were not fully recovered by BL addition^29^. Therefore, we analyzed in detail the *dwf4* root growth and morphology under exogenous BL conditions. Because of the dose-dependent plant growth responses to BRs^9^, we tested the effect of different increasing concentrations of BL on roots of 6-day-old wild type and *dwf4* plants. All BL concentrations tested, from the lowest (10 pM) to the highest (250 pM), had a clear positive effect on the root growth after 1 day for both genotypes when compared to the mock controls (Fig. 4a and Supplementary Fig. 5a). After 2 days, the effect of exogenous BL became detrimental for root growth in Col-0 at concentrations of 100 pM and higher, whereas the growth rates of BL-treated *dwf4* roots were lower than those of the mock-treated wild type at all BL concentrations (Fig. 4A and Supplementary Fig. 5a). Our results support previous observations that the growth rates of BR-deficient *dwf4* roots cannot be fully recovered to the wild type levels by exogenous BL^29^.

**Fig. 4.**
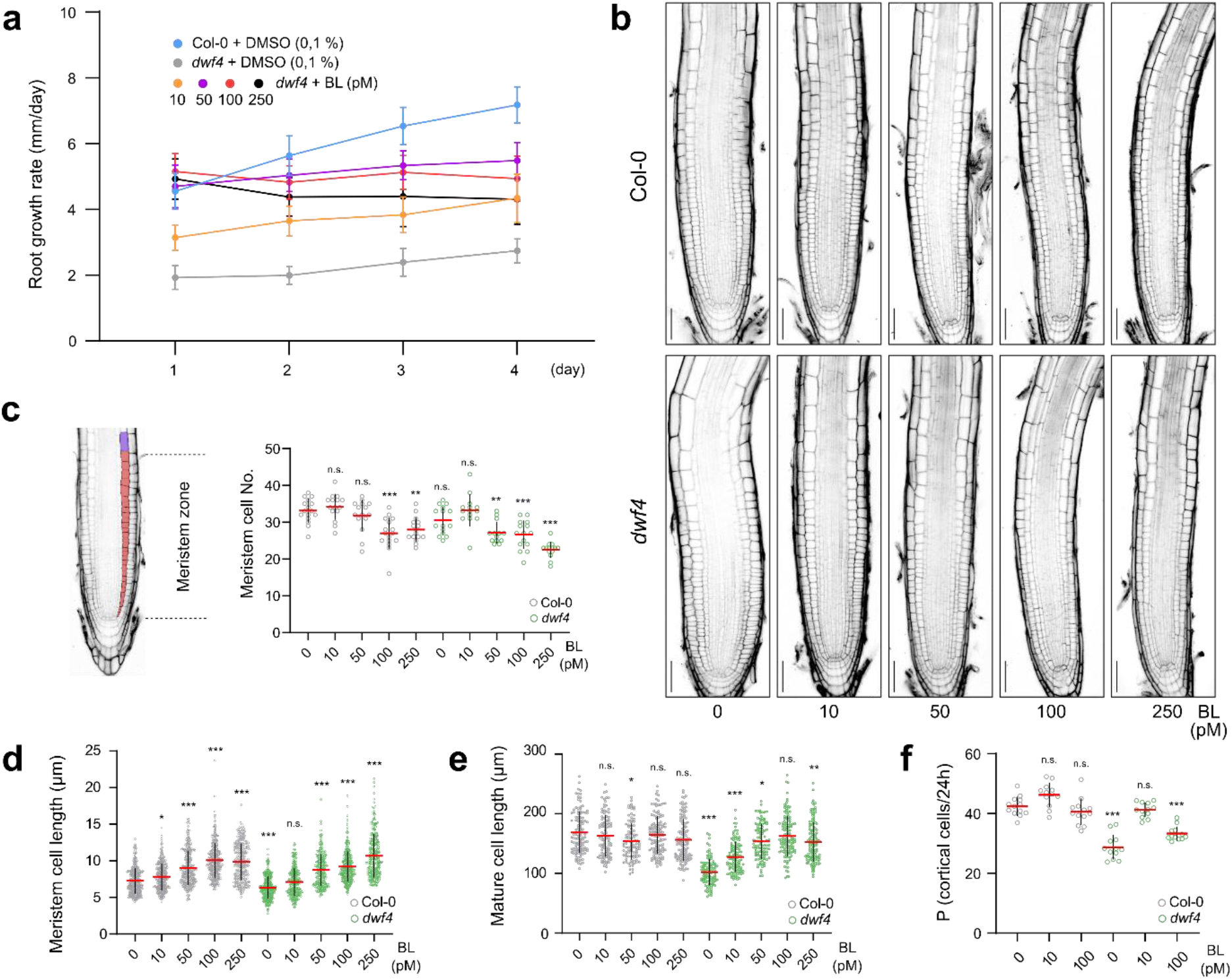
Root meristem and elongation zones have different demands for BRs. **a,** Root growth rates of *dwf4* plants grown on increasing concentrations of brassinolide (BL) compared to Col-0 grown on mock. Six-day-old seedlings were transferred to fresh media containing BL or mock and root growth was followed for 4 days. Values represent the mean ± SD of the root growth rates. *n* ≥ 24 roots analyzed. **b,** Root meristem architecture of Col-0 and *dwf4* plants treated with different concentrations of BL for 24 h. Roots were stained with propidium iodide. Scale bars, 50 μm. **c-e,** Quantification of the cortical meristem cell number (**c**), the cortical meristem cell length, i.e., first 20 cells of individual roots (**d**) and the mature cortical cell length (**e**). Roots were treated with BL for 24 h. *n* ≥ 13 roots analyzed. **f,** Quantification of the root meristem production rate (P) for cortical cells. *n* ≥ 12 roots analyzed. (**c-f**) Circles, red wide bars and black whiskers in the plots represent individual values, means and SD, respectively. The significant differences between Col-0 and the *dwf4* seedlings treated with BL and Col-0 grown on mock (0) were determined by two-way analysis of variance (ANOVA) and Tukey’s multiple comparisons tests. *** *P* < 0.001, ** *P* < 0.01 and * *P* < 0.05.

Next, we examined the morphology of the root meristems of BL-treated Col-0 and *dwf4* plants and found clear differences between roots grown on mock and hormone-supplemented media for 24 h (Fig. 4b). Low hormone concentrations had no significant effect on the meristem cell number, but the high ones, starting from 100 pM, caused a decrease in cell number in both wild type and mutant (Fig. 4c). In addition, BL treatment reduced the root meristem diameter of mutant and Col-0 plants (Supplementary Fig. 5b), whereas the root meristem size was not significantly affected, except when *dwf4* roots were treated with 250 pM BL (Supplementary Fig. 5c). Notably, exogenous BL increased the meristem cell lengths in the two genotypes (Fig. 4d). Interestingly, when treated with 10 pM BL, the *dwf4* roots displayed meristem architecture parameters almost identical to those of the mock-grown wild type. In contrast, the final size of the mature cortical cell of the *dwf4* roots, which was fully recovered by treatment with 100 pM BL, was only partially restored by 10 pM BL (Fig. 4e). Finally, we functionally characterized the root meristem after BL treatments by calculating the number of cells produced in the meristem per unit of time^36^. Root meristem cell production rates of *dwf4* mutant were rescued back to the wild type levels when treated with 10 pM BL, but not with 100 pM (Fig. 4f and Supplementary Fig. 5d,e).

These data imply that meristem and elongation zones require different concentrations of BRs for optimal growth. Whereas low hormone concentrations can restore the *dwf4* meristem architecture, cell length and cell production rates, they are not sufficient to unlock the full elongation potential of the root cells, the reason for the partial root growth recovery of the mutant when grown on 10 pM BL. Conversely, high concentrations of exogenous BRs enable full elongation of the cells in the elongation zone of the mutant root^29^, but simultaneously lead to excessive elongation of meristematic cells, decrease in the meristem cell number and cell production rates and, ultimately reduced root growth.

### DWF4 expression restricts BR production in the *Arabidopsis* root

If low BR levels were retained in the root meristem by maintaining a lessened expression of BR biosynthetic enzyme, then the overexpression of these enzymes should mimic growth under exogenous BL and perturb the meristem architecture in a similar fashion. To test this hypothesis, we expressed the rate-limiting enzyme DWF4 fused to mCHERRY under the control of the constitutive *35S* promoter. As reported previously^37^, DWF4-mCHERRY overexpressing (OE) plants (DWF4-OE) exhibited hypocotyls longer than Col-0 and a range of root phenotypes, including roots with length similar to the wild type, slightly shorter roots and occasionally short, wavy and extremely curled roots, resembling roots treated with high BL concentrations (1 nM) (Fig. 5a). Next, we characterized the BR signaling of DWF4-OE seedlings with long roots. As expected, in roots of these plants, most of BR responsive genes corroborated elevated BR signaling (Fig. 5b). In addition, the phosphorylation status of the BES1 transcription factor in DWF4-OE roots exhibited a shift towards a de-phosphorylated pool, a clear indicator of BR signaling activation (Fig. 5c and Supplementary Fig. 6a). The root meristem architecture of DWF4-OE plants with long roots (Fig. 5d) was similar to that of wild type Col-0 roots treated with a lower (100 pM) BL concentration (Fig. 4) displaying fewer but excessively elongated meristematic cells and smaller root diameter (Fig. 5e-g).

**Fig. 5.**
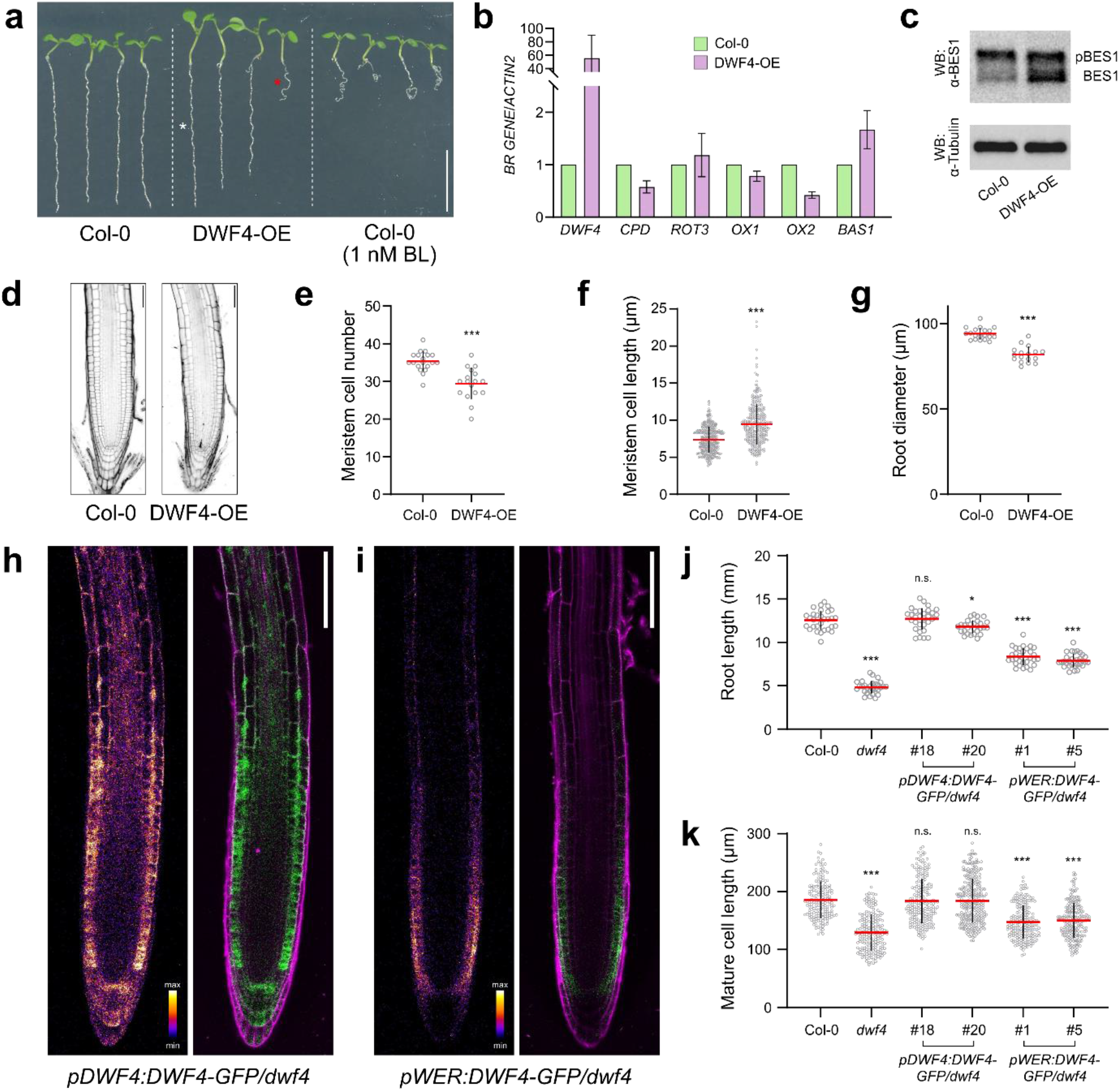
Local DWF4 Expression Determines BR Levels. **a,** Comparison of root phenotypes of 6-day-old seedlings carrying the*p35S:DWF4-mCHERRY* construct [DWF4-overexpression (OE)] with Col-0 seedling grown on mock or media supplemented with 1 nM brassinolide (BL). DWF4-OE plants display long roots (white asterisk) with length comparable to that of Col-0, as well as short ones (red asterisk) comparable to that of roots treated with high concentrations of BL. **b,** Expression of BR-responsive genes in DWF4-OE roots of 6-day-old seedlings measured by real-time quantitative reverse transcription-PCR. Material was collected from seedlings with long roots for DWF4-OE. Expression of *DWF4, CPD, ROT3, BR6OX1* (*OX1*), *BR6OX2* (*OX2*) and *BAS1* was tested. Transcript levels were normalized to the *ACTIN2* gene expression. Three biological replicates were quantified. Bars represent SD. **c,** BES1 phosphorylation status tested by western blot with the α-BES1 antibody in roots of Col-0 and DWF4-OE 6-day-old seedlings. Material was collected from seedlings with long roots for DWF4-OE. Tubulin was used as a loading control. **d,** Root meristem architecture of 6-day-old *Arabidopsis* wild type (Col-0) and DWF4-OE seedlings with long roots. **e-g,** Quantification of different root meristem parameters of 6-day-old Col-0 and DWF4-OE seedlings with long roots. (**e**) Cortical meristem cell number, (**f**) cortical meristem cell length - first 20 cells of individual roots and (**g**) root diameter. Significant differences to Col-0 were determined by Student’s *t* test. *** *P* < 0.001, ** *P* < 0.01 and * *P* < 0.05. *n* ⩾ 17. **h**, Six-day-old *dwf4* mutant roots expressing DWF4-GFP under the control of its native promoter. **i,** Six-day-old *dwf4* mutant roots expressing DWF4-GFP under the control of the *WEREWOLF (WER)* promoter. Note the decrease in GFP signal in the elongation zone. Identical confocal settings were used for (**h**) and (**i**). **j, k,** Quantification of the root lengths (**j**) and mature cortical cell lengths (**k**) of the Col-0, *dwf4* and two independent *dwf4* mutant lines expressing DWF4-GFP under the control of the *DWF4* and *WER* promoters. Six-day-old plants were transferred to fresh medium, lengths were measured after 2 days, and imaged by confocal microscopy. The significant differences between the transgenic lines and the Col-0 control were determined by one-way analysis of variance (ANOVA) and Dunnett’s multiple comparisons tests*** *P* < 0.001, **, *P* < 0.01 and * *P* < 0.05. *n* ⩾ 18 roots analyzed. Roots were stained with propidium iodide. Grey circles, red wide bars and black whiskers in the plots represent individual values, means and SD, respectively. Scale bars represent 1 cm in (**a**) and 50 μm in (**d**, **h** and **i**).

Next, we tested a scenario in which the DWF4 expression was confined to the root meristem zone to assess whether the BR biosynthetic enzyme expression was coupled with hormone production. To this end, we expressed DWF4-GFP under the control of the *WEREWOLF* (*WER*) promoter^38^ that, like the *DWF4* promoter, is active in epidermal meristematic cells. However, whereas the *DWF4* expression level increased and its domain expanded in the elongation zone (Fig. 5h), the *WER* expression remained limited to the epidermal cells and gradually weakened as the cells start to elongate (Fig. 5i). Mutant *dwf4* plants carrying the *pWER:DWF4-GFP* construct had only a partially complemented root length (Fig. 5j and Extended Data Fig. 6b), due to the imperfect rescue of the mature cortex cell length (Fig. 5k). As meristems of roots expressing *pWER:DWF4-GFP* displayed characteristics of normal or roots treated with low amounts of BL (Supplementary Fig. 6c-f), we concluded that their reduced length reflected solely the lack of sufficient DWF4 expression and possible BR biosynthesis in the root elongation zone.

Altogether, our findings reveal that the spatially limited DWF4 expression can restrict the BR production in the *Arabidopsis* root.

## DISCUSSION

Root growth is driven by the coordination of cell division and longitudinal expansion, maintaining the optimal size of the meristematic cell pool and allowing simultaneously cell elongation^26^. Among other factors, BR signaling regulates both cell division and expansion and shows a gradient along the longitudinal root axis^29^. A role for BR biosynthesis in establishing this gradient has been ruled out because of the limited mobility of the hormones and the general belief that every tissue has the capacity to synthesize BRs^20^, albeit the lack of information on their sites of biosynthesis and accumulation.

To explore the importance of BR biosynthesis and distribution for the root growth, we examined the localization of all known enzymes in the pathway. Strikingly, most of the BR biosynthetic enzymes exhibited restricted expression domains especially in the root meristem, implying that BR intermediates have to move over short distances. The mobile nature of BRs had also been indicated by the observation that root meristem defects of the *bri1* receptor mutant were largely rescued by the ectopic expression of BRI1 in the epidermis^32^, a tissue that is separated from the CPD expression domain by several cell layers. Our findings, using the ectopic expression of CPD and BAS1 in the endodermis, experimentally confirm previous suggestions that BRs could move locally^39,40^. In this scenario, locally synthetized BR precursors are able to move within certain root domains to allow the pathway completion and the production of the bioactive compounds. In our experimental setup we could not discriminate whether the precursors or bioactive hormones were able to move and allow the compounds to act in paracrine manner^41^. Simultaneously, the rescue of the *dwf4* mutant root meristems expressing DWF4-GFP under the *WER* promoter, but not the mature cell size, implies that the hormones have a limited mobility over long distances.

Another striking observation was the elevated expression of the BR biosynthetic enzymes in the root elongation zone, correlating with the previously reported BR signaling maximum^29^. Given the inability of BRs to move over long distances^20^, a local increase in hormone synthesis in tissues with a high demand for cell elongation, such as the root elongation zone, would be expected. Links between the sites of BR biosynthetic genes expression and those of active hormone biosynthesis have been proposed previously^42^. In addtion, the local increase in BR biosynthetic gene expression coincides with BR signaling maximum at roughly 200-400 μm from the QC, a region where cells cease to divide and enter the elongation zone^26^. This increase is achieved through either a simple rise in expression levels, such as for CPD, or an expression domain expansion to additional cell files as observed for the DWF4, ROT3 and BR6OX1 enzymes. The question that remains is whether the local increase in biosynthetic gene expression and hormone levels occurs concomitantly. The overexpression of the rate-limiting enzyme DWF4 demonstrates that elevated gene expression leads to increased BR signaling that, in turn, reduces the meristem cell number and negatively affects the root growth, similarly to exogenous BL. Moreover, restriction of DWF4 expression to the meristem region by its expression under the control of *WER* promoter rescues the meristem architecture of the *dwf4* mutant, but not the mature cell length, thus clearly coupling hormone production with local biosynthetic gene expression. Even in the case of auxin, a hormone well known for its directional transport, the localized expression of the auxin biosynthetic enzyme, TRYPTOPHAN AMINOTRANSFERASE OF ARABIDOPSIS1 (TAA1) plays an important role in the formation of local hormone maxima^43^. Therefore, we postulate that the expression maxima of the BR biosynthetic enzymes reflect the hormonal maxima as well, finally determining the local BR signaling response levels. This type of localized biosynthesis places BRs in a unique position among the other major groups of plant hormones which are highly mobile compounds and where transport over long distances often plays crucial role in establishing local hormone maxima^44^.

Collectively, our data demonstrate that root meristem and elongation zones require different BR concentrations for optimal growth. Such dose-dependent responses of roots to exogenous BRs had been shown previously^27,29^. Despite the fact that exogenous BRs, even when applied at extremely high concentrations, have no negative impact on the final size of the mature cells in the root elongation zone, the root growth is strongly impaired^29^. However, increased BR levels have negative effect on meristem activity^27^. Root growth defects caused by elevated BRs can be explained with a Sizer mechanism for cell elongation and differentiation during root growth^45^, which predicts that cells sense their own length and cease to expand after a threshold value is reached. According to this model, the final length of differentiated cells is independent of changes in the meristematic activities and therefore it can explain the retarded root growth phenotypes observed in situation with unaltered mature cell length and decreased meristematic activity. Because minute quantities of exogenous BL can restore the root meristem architecture and function of the *dwf4* mutant but not the elongation zone defects, we postulate that a low level of BR signaling is kept in the root meristem by maintaining low amounts of endogenous BRs. BRs. Hence, we propose a model (Fig. 6a) in which the BR signaling output reflects the existence of a hormone gradient along the longitudinal root axis. In BR deficient mutants, the elongation of all root cells is impaired, causing two distinct phenotypic defects, a decreased final cell length and decreased meristematic activity. Both phenotypic defects can be fully rescued by exogenous BL, but not simultaneously. Whereas low BL concentrations are able to rescue the root meristem architecture of the biosynthetic mutants, only high concentrations enable full cell elongation. Nevertheless, high-concentration treatments lead to untimely elongation of meristematic cells and to a decrease in meristem cell numbers and mitotic activity cell production rate (Fig. 6b).

**Fig. 6.**
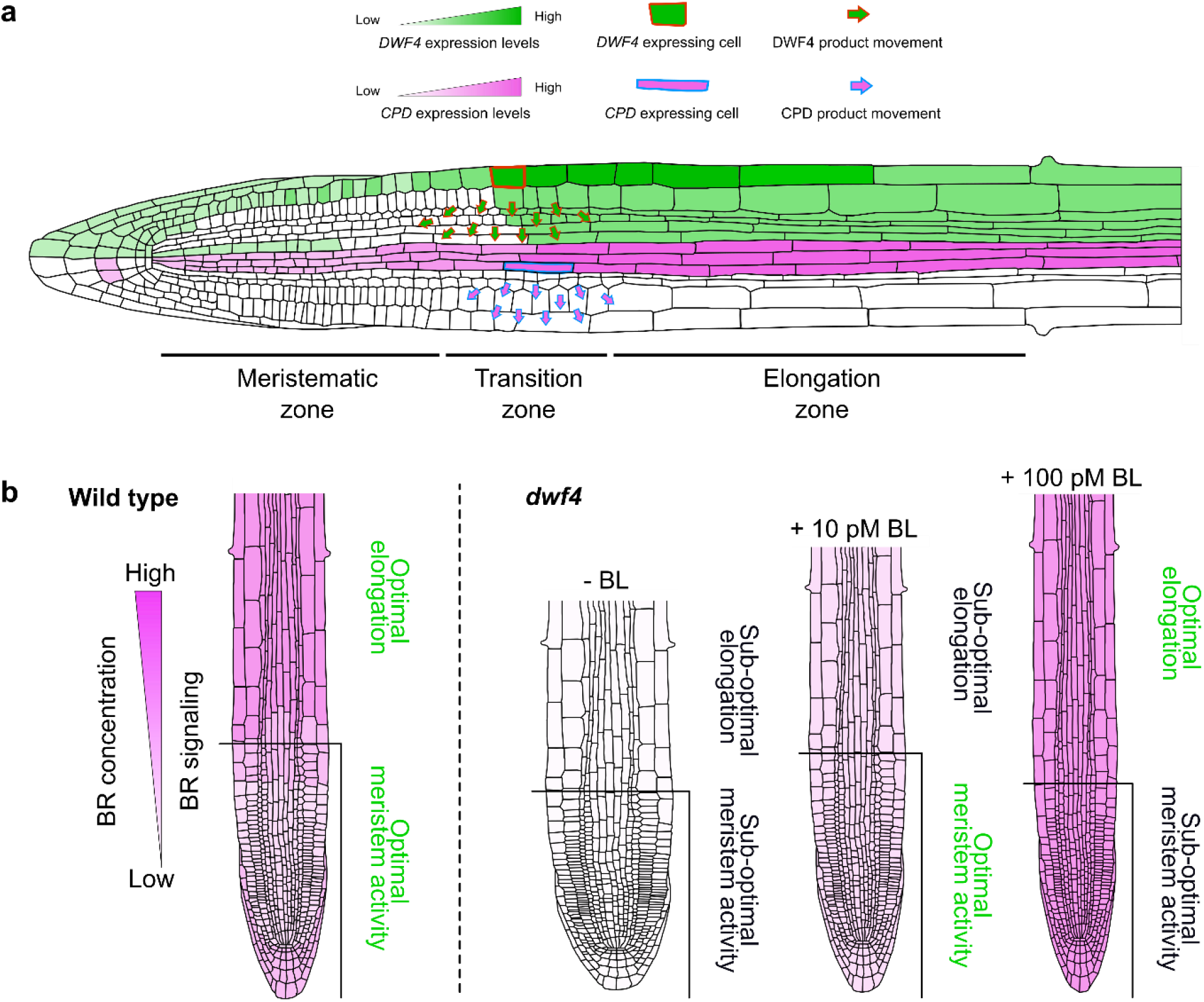
Localized BR biosynthesis coordinates root growth. **a**, Not always overlapping expression domains of BR biosynthetic enzymes. Expression patterns DWF4 and CPD, two consecutive enzymes in the biosynthetic pathway, are shown. BR intermediates need to be exchanged between neighboring cells to be converted into bioactive molecules. This movement is restricted and creates local BR maxima and a hormone concentration gradient along the longitudinal root axis. **b,** BR signaling shows a gradient along the longitudinal root axis and enables optimal cell division rates in meristem and cell expansion in the elongation zone. This signaling gradient corresponds to local expression maxima of the BR biosynthetic genes and, hence, local BR levels. In *dwf4*-biosynthetic mutant roots, meristematic cells divide at a reduced rate and their elongation is impaired, resulting in a short root phenotype. When *dwf4* roots are treated with low concentrations of BL, the meristem activity is recovered, but not the cell elongation. Increased concentrations recover the *dwf4* cell elongation defects, but simultaneously deplete cells from the meristems. Therefore, both treatments lead to *dwf4* roots shorter than those of the wild type, demonstrating that low BR levels in the meristems and high BR levels in the elongation zone are needed for optimal root growth.

In summary, we postulate that the BR signaling levels correspond to local hormonal accumulation that is achieved through regulation of the expression of the BR biosynthetic enzymes and by a radial short-distance transport of active BRs and their precursors. Due to the limited mobility, local BR production leads to local hormone accumulation, which, in turn, triggers signaling and enables timely transition of meristematic cells to the elongation zone.

## Methods

### Plant material

*Arabidopsis thaliana* (L.) Heynh. Col-0 plants were used for all experiments. For the phenotypic analysis, following mutant lines were used: *dwf4-102* (At3g50660)^46^, *cpd* (At5g05690)^3^, *det2-1* (At2g38050)^47^, *rot3* (At4g36380)^48^, *cyp90d1* (At3g13730)^49^, *br6ox1-1* (At5g38970)^50^, *br6ox2-2* (At3g30180)^50^ and *bri1/brl1/brl3* (At4g39400 / At1g55610 / At3g13380)^51^. Mutants *rot3* and *cyp90d1*, and *br6ox1* and *br6ox2* were crossed to obtain double mutants. Genotyping primers are listed in Supplementary Table 1. The transgenic *Arabidopsis* lines expressing *pDET2:DET2-GFP-GUS*^52^ and *pBES1:BES1-GFP*^1^ have been described previously.

### Growth conditions

Seeds were surface-sterilized with the sterilization buffer (80% [v/v] ethanol, 20% [v/v] sodium hypochlorite), stratified for 2 days in the dark at 4°C, and grown vertically on half-strength Murashige and Skoog (½MS) agar plates, supplemented with 1% (w/v) sucrose at 22°C, with a 16-h/8-h light/dark photoperiod. For the phenotypic analysis 6-day-old seedlings were transferred to fresh plates for 3 additional days for root growth measurements or transferred to soil and imaged after 3 weeks.

### Treatments and root growth assays

For the BL treatments of the *dwf4* mutants, plants were grown on agar plates for 6 days until segregating homozygous mutants were recognizable (wild type-looking seedlings consisting of Col-0 and *dwf4* heterozygous plants were used as a wild type control), whereafter the seedlings were transferred to fresh media containing different BL concentrations (OlChemIm Ltd.). BL was kept at different stock concentrations in DMSO and was diluted 1000× to reach the final concentrations in the media. For the mock treatment, DMSO was at a final concentration of 0.1% (v/v). Root growth was recorded every 24 h for 4 days. For confocal microscope observations, the procedure was repeated, root tips were stained with propidium iodide (Sigma-Aldrich) and imaged 24 h post-treatment. Meristem area size was determined as a region starting from QC to the first elongating cell (length of the cell is two times greater than width) and meristem cell number was determined as a number of cortical cells in single file spanning from QC to the first elongated cell. To measure length of mature cortical cells, cells from root region where root hairs start to emerge (a sign of cell differentiation after the elongation ceased) were imaged. Root meristem production rate (P) was measured according to^36^. Six-day-old Col-0 and *dwf4* seedlings were transferred to fresh agar plates containing different BL concentrations and grown for 3 days. On the third day, root tips were marked and grown for additional 24 h hours. Root elongation (E) of individual roots was recorded, roots were stained with propidium iodide and imaged with confocal microscope. For each root, length of at least 30 mature cortical cells was measured and mean cell length was calculated (L_avg_). Meristem production rate (P) was calculated using the equation P=E/ L_avg_.

### Microscopy and image analysis

Seedlings were imaged on a Leica SP8X confocal microscope. GFP was excited at 488 nm and acquired at 500 to 530 nm. mCHERRY was excited at 594 nm and acquired at 600 to 650 nm. For meristem imaging, roots were stained with propidium iodide (PI) (Sigma-Aldrich). Images were taken by 20× or 40× objectives. Vertical imaging was done using a Carl Zeiss Axio Observer.7 armed with a Visiscope spinning disk confocal unit based on Yokogawa CSU-W1-T2 equipped with a VS-HOM1000 excitation light homogenizer and a PRIME-95B Back-Illuminated sCMOS camera, and a Plan-Apochromat 20x/0.8 M27 objective. For time-lapse microscopy experiments with plants expressing *pDWF4:DWF4-GFP, pCPD:CPD-GFP, pROT3:ROT3-GFP* and *pBES1:BES1-GFP*, 6-day-old seedlings were transferred to a 3D-printed triple microscopy chamber (5 seedlings to each) on ½MS agar blocks, supplemented with propidium iodide (PI) and imaged overnight at 25-min intervals. Time series were assembled with stitching of tiled 3D microscopic image acquisitions^53^ and subsequently analyzed with Fiji or LAS X software.

### Generation of constructs and transgenic lines

To generate transcriptional reporter lines of BR biosynthetic enzymes, promoter regions roughly 2 kbp upstream from the start codons of *DWF4*, *CPD*, *ROT3* and *CYP90D1* were amplified by PCR and cloned into pDONRP4-P1R by means of the Gateway BP clonase enzyme mix (Thermo Fisher Scientific). The *BR6OX1* promoter was amplified and cloned with the restriction enzymes *Bam*HI and *Xho*I into pENTRL4-R1^54^. For *BR6OX2*, a 6.5-kbp promoter region was incorporated into pDONRP4-P1R by assembling four PCR fragments with the Gibson Assembly technology (New England BioLabs). The protocol for the Gibson assembly, including the primer design, was according to the instruction manual of the Gibson Assembly Master Mix. All promoters were recombined into pMK7S*NFm14GW,0^55^ with the Gateway LR clonase enzyme mix (Thermo Fisher Scientific) to generate the expression constructs *PROMOTER:NLS-GFP*, each carrying a kanamycin resistance gene for plant selection. For the *pROT3:NLS-mCHERRY* construct, the promoter-carrying entry clone was combined with entry clones carrying the NLS signal peptide and the *mCHERRY* gene. These constructs were introduced into the Col-0 wild type by means of the floral-dip method^56^. The *pROT3:NLS-mCHERRY* and *pCPD:NLS-GFP* lines were crossed to obtain a co-expressing line.

To generate translational reporter lines of the BR-biosynthetic enzymes, the coding sequences of the full genomic fragments of *DWF4, CPD, CYP90D1, BR6OX1* and *BR6OX2* were amplified by PCR and cloned by BP clonase enzyme mix into pDONR221 (Thermo Fisher Scientific). The genomic DNA fragment of *ROT3* was isolated by overlap extension PCR and cloned into pDONR221. For *GFP-CYP85A2*, the signal peptide sequence (first 78 bp starting from ATG)^57^ was included into the GFP cloning primer and *SP-GFP* was PCR amplified and introduced into pDONR221. The genomic sequence of *CYP85A2* without the first 78 bp was cloned into pDONR2R-P3. Entry clones were combined with destination vectors in LR reactions to obtain expression clones with GFP-tagged enzymes (pB7m34GW for *DWF4, ROT3*, *CYP90D1*, *BR6OX1* and *BR6OX2* and pK7m34GW for *CPD*). The expression clones were introduced into corresponding mutant backgrounds with the floral dip method. To generate the*pWER:DWF4-GFP* construct, the cDNA of DWF4 was amplified by PCR, cloned into pDONR221 and subsequently recombined with the *WER* promoter-carrying pDONRP4-P1R^58^ into the pH8m34GW-FAST destination vector. The DWF4-OE line was constructed by recombination of pDONR221 with genomic *DWF4* into pDONRP4-P1R-35S (cauliflower mosaic virus *(CaMV)35S* promoter) and of pDNORP2R-P3 into the pB7m34GW destination vector. For the *CPD* and *BAS1* ectopic expression lines, cDNA was cloned into pDONR221 and recombined in an LR reaction with promoter-carrying entry clones, *SCR* for *CPD* and *RPS5A* and *SCR-XVE*^59^ for *BAS1*. *CPD-mCHERRY* and *BAS1-GFP* were recombined into the pB7m34GW and pH8m34GW-FAST vectors, respectively. Cloning primers are listed in Supplementary Table 1.

### Western blot analysis

For the BES1 Western Blot analysis, experiments were done in triplicate. For protein extraction, approximately 50 roots of 6-day-old seedlings were collected. For the DWF4-OE line, roots of all sizes were pulled together. Plant material was frozen in liquid nitrogen, grinded by Retsch MM400, and homogenized in 100 μl ice-cold homogenization buffer (1% [v/v] SDS, 25 mM Tris/HCl, pH 7.5, 150 mM NaCl, 10 mM DTT and a Roche Complete protease inhibitor 1 tablet/10 ml), put to the ice for 30 min. The homogenates were centrifuged twice (10 min, 14,000 rpm) at 4°C. After addition of LDS [4×] and Sample Reducing Agent [10×], the samples were heated for 10 min at 70°C, centrifuged again, separated on 4-15% [v/v] SDS-PAGE stain-free protein gel (Bio-Rad Laboratories), and blotted on Trans-Blot^®^ Turbo™ Mini PVDF Transfer Packs. Membranes were blocked at 4°C in 5% [v/v] Difco™ Skim Milk. For immunodetection, anti-BES1 antibody at 1:5000 was used as primary antibody and donkey anti-Rabbit (Merck) at 1:10000 was used as secondary antibody. For tubulin detection anti-Tubulin (Abcam) at 1:5000 was used as primary antibody and sheep anti-Mouse (Merck) at 1:10000 was used as the secondary antibody. The proteins were detected by ChemiDoc™ MP Imaging System (Bio-Rad Laboratories). For the BES1 dephosphorylation assay, the ratio of the dephosphorylated BES1 to the total BES1 proteins was quantified according to the signal intensity. The loading was adjusted to an equal level based on the amount of tubulin. Signal intensities were determined with Image Lab (Bio-Rad Laboratories).

### Quantitative RT–PCR

Total RNA was extracted by quantitative RT-PCR from Col-0 and DWF4-OE seedlings 6 days post germination with the RNeasy Mini Kit (Qiagen). For DWF-OE, all root types (short and long) were collected and pooled together. Genomic DNA was eliminated by on-column digestion with RQ1 RNase-free DNase (Promega) during the isolation procedure. cDNA was generated from 1 μg total RNA with qScript cDNA SuperMix (Quantabio) and analyzed on a LightCycler 480 II apparatus (Roche) with the SYBR Green I Master mix (Roche) according to the manufacturer’s instructions. Expression levels were normalized to those of *ACTIN2*. Primers are listed in Supplementary Table 1.

### Quantification and statistical analysis

All statistical analyses were done in GraphPad Prism 8 software. Significance of differences was determined with two-tailed Student’s unpaired *t*-test analysis for binary comparisons. Comparisons of more than two genotypes was done by one-way ANOVA and Dunnett’s multiple comparisons test was subsequently used in the comparison procedure. For the *dwf4* mutant rescue treatments with BL, a two-way ANOVA was utilized and Tukey’s multiple comparison test was subsequently used in the comparison procedure. Asterisks illustrate the *P* value: *P* < 0.001 is ***, *P* < 0.01 is ** and *P* < 0.05 is *.

#### Reporting Summary

Further information on research design is available in the Nature Research Reporting Summary linked to this article.

#### Data availability

All supporting data are included in the Supplementary Information. Source data are provided with this paper.

## Acknowledgments

We thank Yanhai Yin for providing the anti-BES1 antibody, Zhaojun Ding, Mitsunori Seo and Csaba Koncz for providing published materials, Tom Beeckman, Jiří Friml, Gerrit Beemster, Roman Pleskot and Klaas Yperman for useful discussions, and Martine De Cock for help in preparing the manuscript. This work was supported by the Research Foundation-Flanders (project G022516N to E.R. and a postdoctoral fellowship 12R7819N to N.V.), the Chinese Scholarship Council (predoctoral fellowships to Y.W. and B.G.), European Research Council (Grant No. 803048 to M.F.) and ERDF project “Plants as a tool for sustainable global development” (No. CZ.02.1.01/0.0/0.0/16_019/0000827 to M.K., P.J. J.O. and M.S.).

## Author contributions

N.V. and E.R. initiated the project. N.V. and Y.W. designed and performed most of the experiments. I.V., M.F., B.G., M.K., P.J. and J.O. performed experiments and analyzed data. N.V. and E.R. wrote the manuscript. All authors revised the manuscript.

## Competing interests

The authors declare no competing interests.

## Additional information

Supplementary information is available for this paper

**Correspondence and requests for materials should be addressed** to N.V. and E.R.

## Notes

### Competing Interest Statement

The authors have declared no competing interest.

