## Supplemental figures and table for "Local brassinosteroid biosynthesis enables optimal root growth"

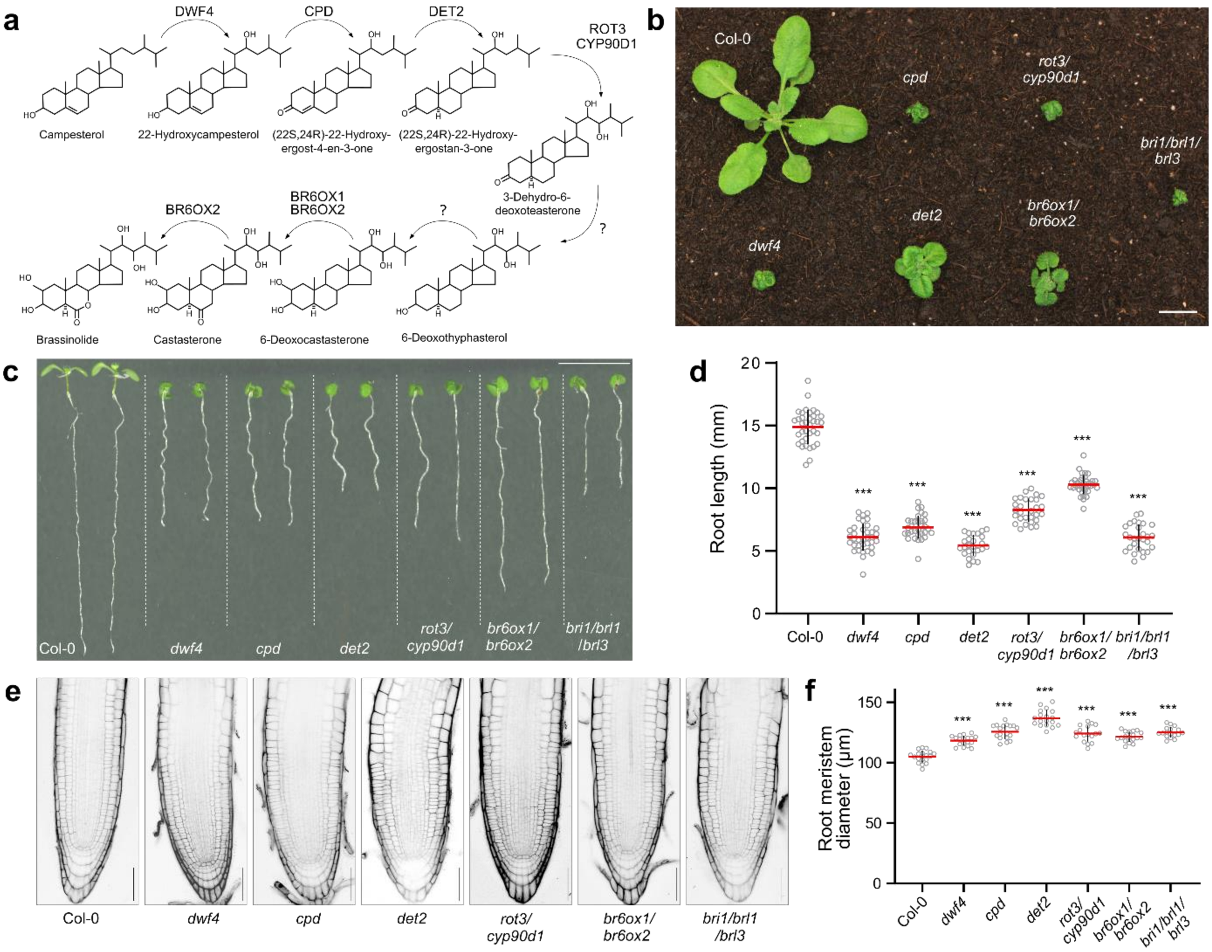


**Supplementary Fig. 1. BR biosynthetic mutants are dwarfs with reduced root growth. a,** Simplified BR biosynthetic pathway with all known enzymes and their presumed positions within the pathway. **b,** Dwarf phenotypes with compact rosettes of the 3-week-old *dwf4*, *cpd*, *det2*, *rot3/cyp90d1* and *br6ox1/br6ox2* mutants defective in BR biosynthetic enzymes and the *bri1/brl1/brl3* triple mutant defective in BR signaling. **c,** Nine-day-old BR biosynthetic and signaling mutants with roots shorter than those of the wild type Col-0 control. **d,** Quantification of the root lengths of BR biosynthetic and signaling mutants shown in (**c**). Six-day-old plants were transferred to fresh medium and root growth was measured after 3 days, $n \geqslant 25$ roots analyzed. **e,** Increased root diameter of BR biosynthetic and signaling mutants. Six-day-old roots were stained with propidium iodide. **f,** Quantification of the root meristem diameter shown in (**e**), $n \geqslant 17$ roots analyzed. Grey circles, red wide bars and black whiskers in (**d**) and (**f**) represent individual values, means and SD, respectively. The significant differences between mutants and the Col-0 control were determined by one-way analysis of variance (ANOVA) and Dunnett's multiple comparisons tests. *** $P < 0.001$, ** $P < 0.01$ and * $P < 0.05$. Scale bars represent 1 cm in (**b**) and (**c**) and 50 µm in (**e**).

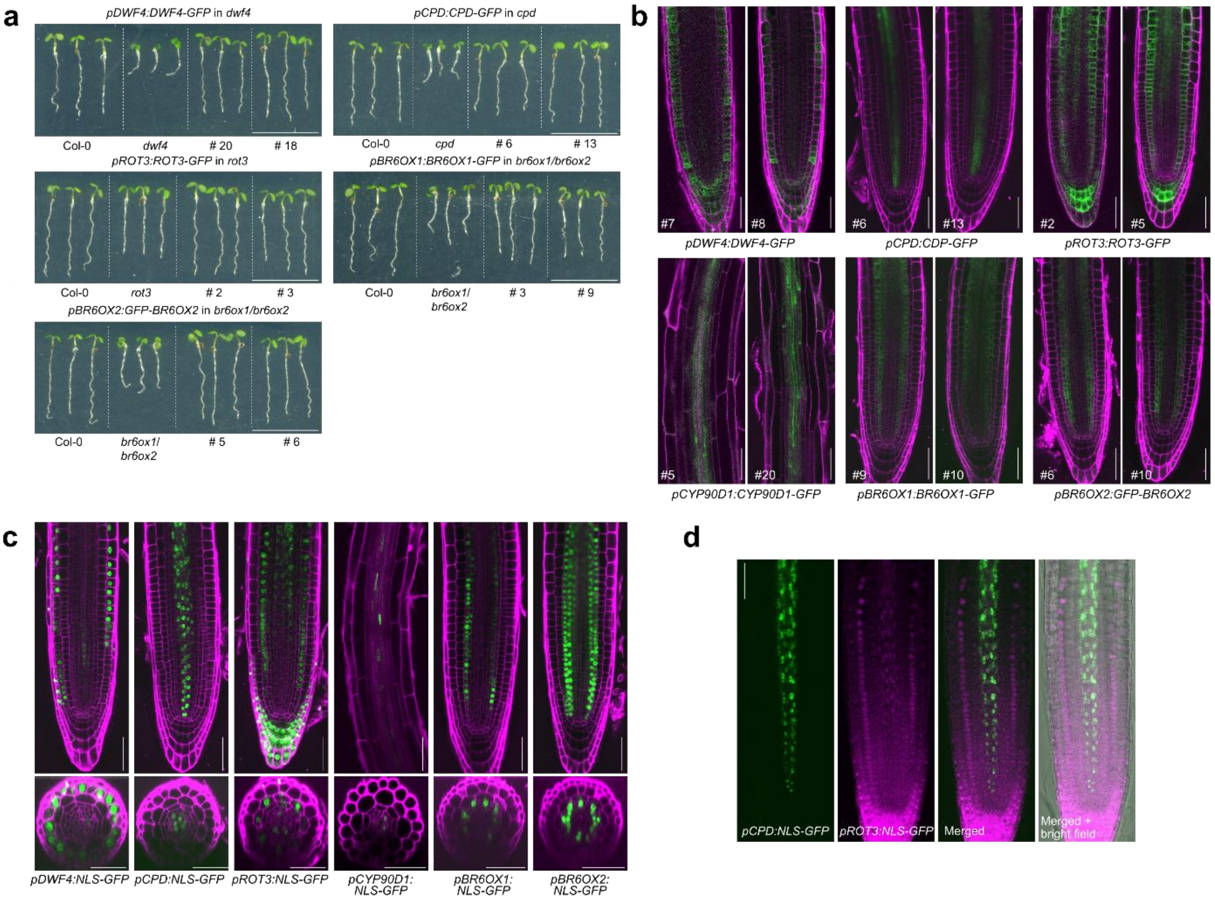


**Supplementary Fig. 2. The GFP-tagged BR biosynthetic enzymes are functional and show similar expression patterns in multiple independent transgenic lines. a,** Rescue of the BR biosynthetic mutants by expression of the corresponding GFP-tagged enzymes under the control of their endogenous promoters. Two independent transgenic lines for each gene are compared with the wild type (Col-0) and the corresponding mutant. Six-day-old seedlings are shown. Scale bars, 1 cm. **b,** Consistent expression patterns of GFP-tagged BR biosynthetic enzymes under the control of their endogenous promoters in multiple independent transgenic lines. Root meristems of the 6-day-old seedlings from two independent transgenic lines for each GFP-tagged enzyme are shown. For seedlings expressing *pCYP90D1:CYP90D1-GFP* only the root elongation zone is shown, since the signal in the root apical meristem is extremely weak. Roots were stained with propidium iodide. Scale bars, 50 µm. **c,** Expression patterns of the *PROMOTER-NLS-GFP* reporters for all BR biosynthetic genes. For each reporter line, 6-day-old root meristems and cross section are shown. Scale bars, 50 µm. **d,** Co-expression of *NLS-GFP* and *NLS-mCHERRY* reporters under the control of *CPD* and *ROT3* promoters, respectively. Scale bar, 50 µm. Roots were stained with propidium iodide.

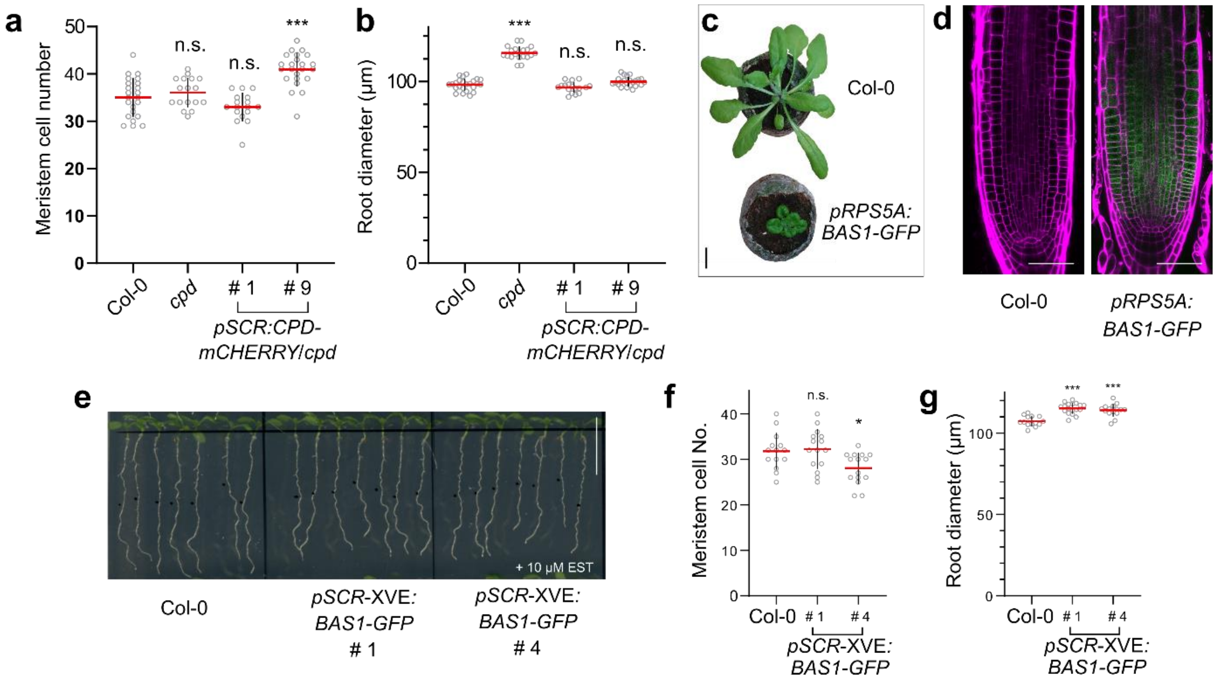


**Supplementary Fig. 3. Ectopic expression of CPD and BAS1 in the *Arabidopsis* root. a,** Quantification of the root meristem cell number of *cpd* seedlings expressing the *pSCR:CPD-mCHERRY* construct. **b,** Reverted root meristem diameter of the *cpd* mutant to that of the wild type upon ectopic expression of *CPD* in the endodermis. Six-day-old seedlings of the two independent transgenic lines were measured, $n \geqslant 17$ roots analyzed. **c,** Active BR depletion by the BAS1-GFP overexpression causing a typical BR deficient phenotype. Scale bar, 1 cm. **d,** Thickening of the meristem and shortening of the meristem cells caused by the BAS1-GFP overexpression, typical for BR-related mutants. Scale bars, 50 µm. **e,** Delayed root growth by the inducible, tissue-specific expression of BAS1-GFP in the endodermis. Two independent transgenic lines were compared with the wild type (Col-0). Four-day-old seedlings were transferred to fresh medium containing 10 µM β-estradiol and imaged after 3 days. Scale bar, 1 cm. **f, g,** Increased root meristem diameter (**f**) and decreased root meristem size (**g**) by the inducible ectopic expression of BAS1-GFP in the root endodermis, $n \geqslant 18$ root analyzed. Grey circles, red wide bars and black whiskers in plots represent individual values, means and SD, respectively. The significant differences between transgenic lines and the Col-0 control were determined by one-way analysis of variance (ANOVA) and Dunnett's multiple comparisons tests. *** $P < 0.001$, ** $P < 0.01$ and * $P < 0.05$.

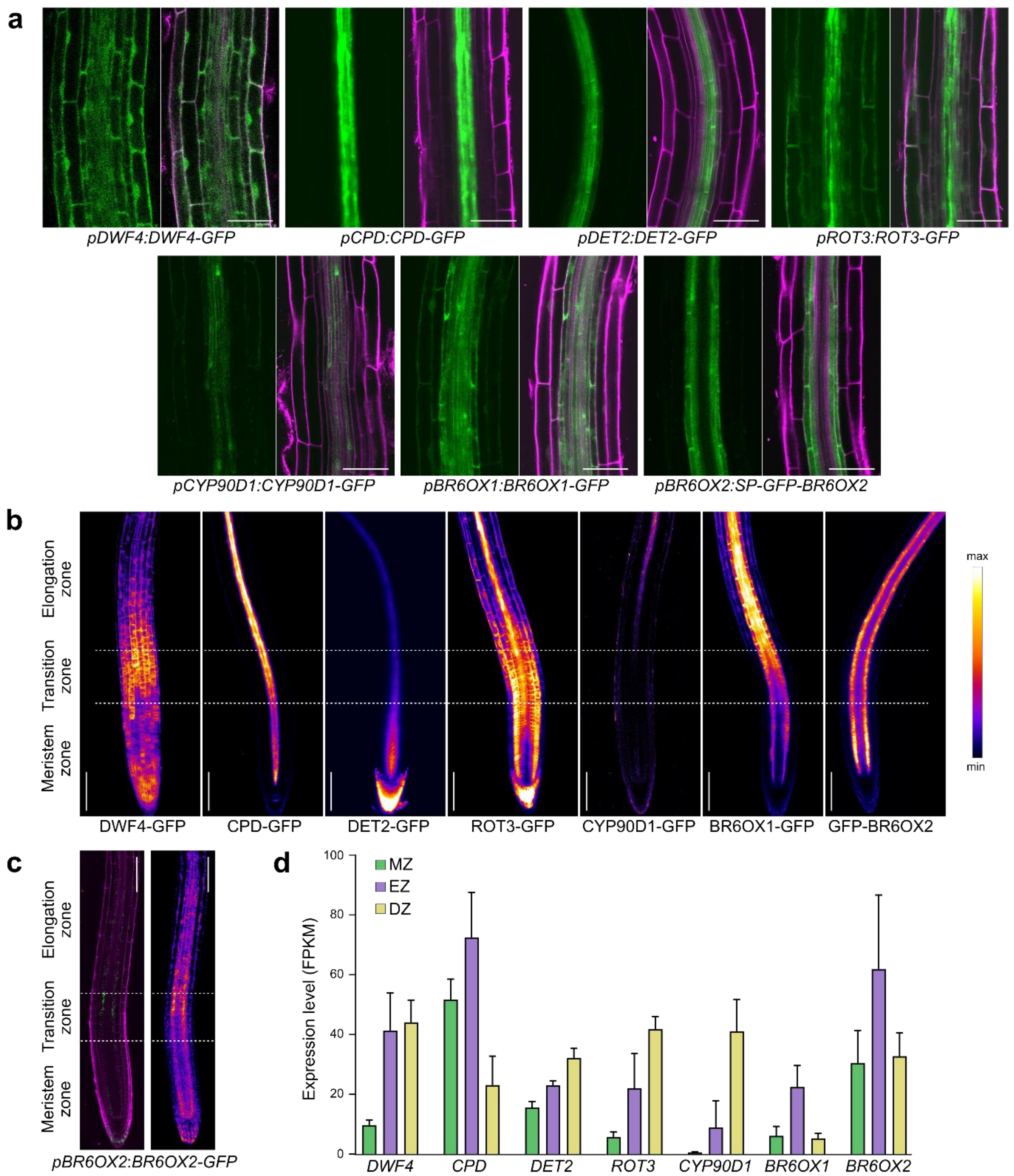


**Supplementary Fig. 4. Expression patterns of BR biosynthetic enzymes in the root elongation zone. a,** The expression domains of several genes expanding in the root elongation zone. DWF4-GFP expression is very broad and the GFP signal can be observed in all cell types of the elongation zone. The CPD-GFP expression is restricted to the stele of the elongation zone. The DET2-GFP expression is limited to the stele, similarly to the expression of CPD-GFP. The ROT3-GFP expression has maxima in the stele, but a signal also occurs in all other cell types. The expression of the ROT3 homolog, CYP90D1, is limited to the stele. BR6OX1 and BR6OX2 both have expression maxima in endodermis and pericycle, but the expression pattern of BR6OX1 is expanded in all cell types. Roots were stained with propidium iodide. Scale bars, 50 µm. **b,** Increase in the expression levels of most of the BR biosynthetic enzymes in the transition and elongation zones as visualized by color-based fluorescence intensity coding. All reporter lines showed an increase in expression higher in the root except for DET2-GFP and GFP-BR6OX2. Maximum Z projections of the same roots from Fig. 2a are shown. Scale bars, 100 µm. **c,** Roots expressing *pBR6OX2:BR6OX2-GFP* (left panel). Increased GFP signal can be observed in elongation zone of as visualized by color-based fluorescence intensity coding (right panel). Scale bars, 100 µm. **d,** Expression levels of BR biosynthetic genes in the *Arabidopsis* root. All genes show an increased expression in the elongation zone. Extracted from (Huang and Schiefelbein, 2015). MZ, meristematic zone; EZ, elongation zone; DZ, differentiation zone. Bars represent SD.

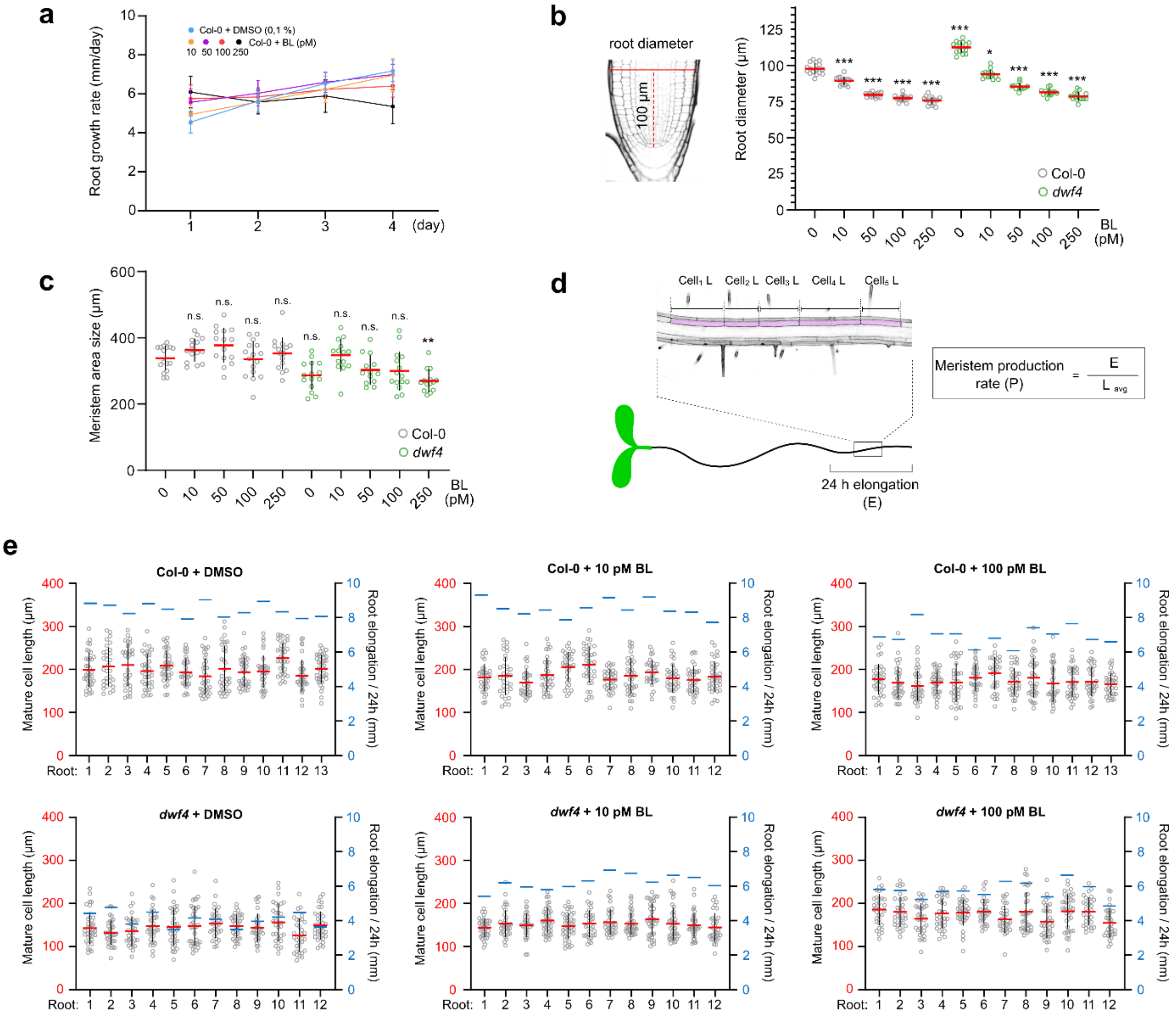


**Supplementary Fig. 5. Effects of exogenous brassinolide (BL) on root growth. a,** Root growth rates of Col-0 plants grown on increasing concentrations of BL. Six-day-old seedlings were transferred to fresh media containing BL or mock and root growth was followed for 4 days. Values represent the mean ± SD of the root growth rates. $n \geq 24$ roots analyzed. **b,** Quantification of the root meristem diameter of BL-treated 6-day-old Col-0 and *dwf4* seedlings. Roots were treated with BL for 24 h, $n \geq 13$ roots analyzed. **c,** Quantification of the root meristem area size measured from the quiescent center to the first elongated cell of Col-0 and *dwf4* treated with increasing concentrations of BL. Roots were treated with BL for 24 h, $n \geq 13$ roots analyzed. The significant differences between Col-0 and the *dwf4* seedlings treated with BL and Col-0 grown on mock (0) in (**b** and **c**) were determined by two-way analysis of variance (ANOVA) and Tukey's multiple comparisons tests. ***$P < 0.001$, ** $P < 0.01$and * $P < 0.05$. **d,** Experimental setup for measurements of the root meristem cell production rate (left panel). Six-day-old Col-0 and *dwf4* seedlings were transferred to agar plates supplemented with BL and mock, and after 3 days of growth, elongation of individual roots was recorded for the period of 24 h (E). Subsequently, roots were imaged with confocal microscope, and average mature cell length was calculated ($L_{avg}$) for each root and used to calculate meristem production rate (P). **e,** Quantification of mature cell length of individual roots used to calculate meristem production rate in Fig. 4f. (**b**, **c**, **e**) Circles, red wide bars and black whiskers in the plots represent individual values, means and SD, respectively. Blue bars in (**e**) represent elongation (E) of individual roots over 24 h.

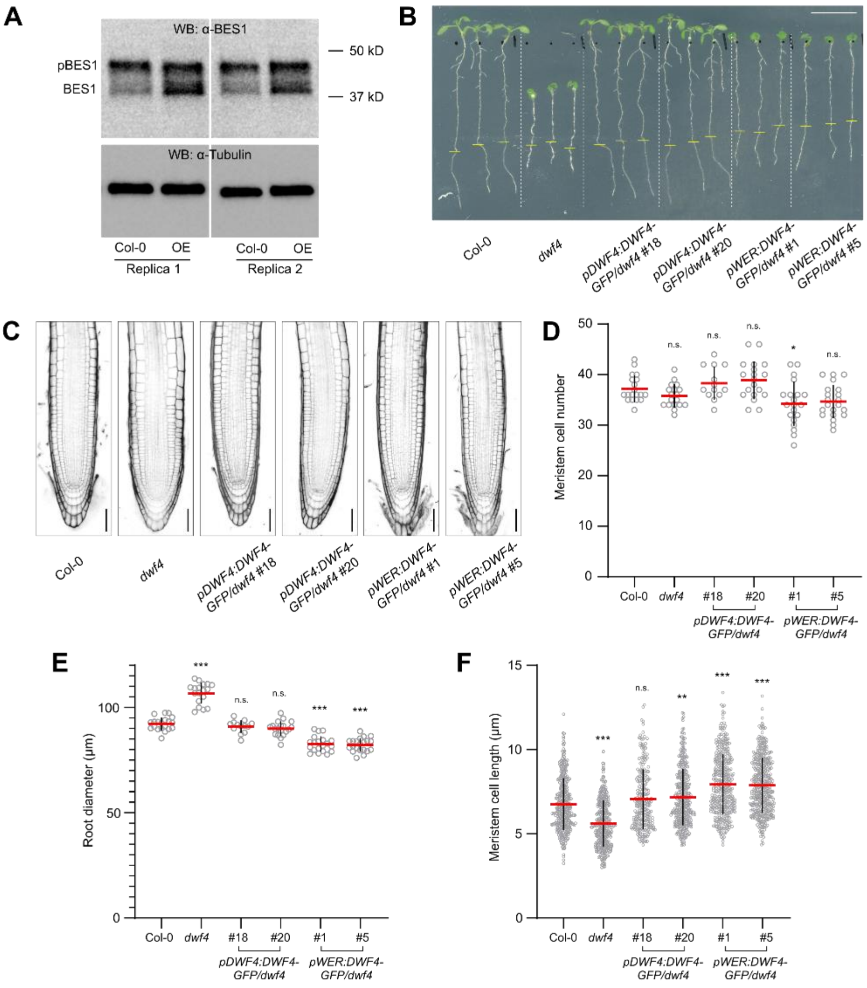


**Supplementary Fig. 6. *DWF4* ectopic expression. a,** Two biological replicates of the western blot shown in Fig. 5c. **b,** Eight-day-old seedlings expressing DWF4-GFP in *dwf4* mutant background under the control of native or *WER* promoter. Two independent transgenic lines for each construct are compared with wild type (Col-0) and *dwf4*. Plants were grown for 6 days until the point where homozygous *dwf4* plants can be recognized and transferred to fresh agar plates. Root tips were marked (yellow bar), and grown for two additional days. **c,** Confocal images of root meristems of Col-0, *dwf4* and two independent *dwf4* mutant lines expressing DWF4-GFP under the control of the *DWF4* and *WER* promoters. Six-day-old plants were imaged. Roots were stained with propidium iodide. Scale bars 50 µm. **d-f,** Quantification of cortical meristem cell number (**d**), root meristem diameter (**e**) and cortical meristem cell length - first 20 cells of individual roots (**f**). $n \geqslant 13$ roots analyzed. Circles, red wide bars and black whiskers in plots represent individual values, means and SD, respectively. The significant differences between the transgenic lines and the Col-0 control were determined by one-way analysis of variance (ANOVA) and Dunnett's multiple comparisons tests. *** $P < 0.001$, ** $P < 0.01$ and * $P < 0.05$.

### Supplementary Table 1 Oligonucleotides used in this study

| Primer name | Primer sequence |
|---|---|
| **Cloning primers** | |
| pDWF4_F | GGGGACAACTTTGTATAGAAAAGTTGTTGTCAACATATACATAATTCCTTACAG |
| pDWF4_R | GGGGACTGCTTTTTTGTACAAACTTGTGGAGCTAGTTTCTCTCTCTCTCTC |
| pCPD_F | GGGGACAACTTTGTATAGAAAAGTTGTTCTAAGTCCAAAAACATATACAATCC |
| pCPD_R | GGGGACTGCTTTTTTGTACAAACTTGTTGAAGAAGAAGATGATGATGAGAGAAG |
| pROT3_F | GGGGACAACTTTGTATAGAAAAGTTGTTCAATAAACACTATGCTCTATGCGATGG |
| pROT3_R | GGGGACTGCTTTTTTGTACAAACTTGTACGTAAACAGTTAAACCGGTTTCTTGGG |
| pCYP90D1_F | GGGGACAACTTTGTATAGAAAAGTTGTTCCAAATGAAAACAACACCACAATTCCC |
| pCYP90D1_R | GGGGACTGCTTTTTTGTACAAACTTGTTAGTGTTTTAACTTAATTAGCCAGAGACG |
| pBR6OX1_F | GGGGACAACTTTGTATAGAAAAGTTGTTAAGGATGCCGTGTGTTTGGAAGTATG |
| pBR6OX1_R | GGGGACTGCTTTTTTGTACAAACTTGTCTCACTCTGTTTTCTGCTCTGTTTTAAAAC |
| pBR6OX2_1_F | CCAAATCCAGACGTTAAATG |
| pBR6OX2_1_R | GAAATTCGATTCTGATTCCG |
| pBR6OX2_2_F | CGGAATCAGAATCGAATTTC |
| pBR6OX2_2_R | CTGCTTCACAAATCTGTAAC |
| pBR6OX2_3_F | GTTACAGATTTGTGAAGCAG |
| pBR6OX2_3_R | CTAATTGCATTGTTAACTTA |
| pBR6OX2_4_F | TAAGTTAACAATGCAATTAG |
| pBR6OX2_4_R | CTTTGTGTGTTTTTTTGCTC |
| Gibson_P4-P1R_F | GAGCAAAAAAACACACAAAGCAAGTTTGTACAAAAAAGTTGAACGAG |
| Gibson_P4-P1R_R | CTCATTTAACGTCTGGATTTGGCAACTTTTCTATACAAAGTTGG |
| pSCR_F | GGGGACAACTTTGTATAGAAAAGTTGTTCTTTTTAAGGTTTTTGTTGCC |
| pSCR_R | GGGGACTGCTTTTTTGTACAAACTTGTGGAGATTGAAGGGTTGTTGG |
| DWF4_F | GGGGACAAGTTTGTACAAAAAAGCAGGCTTTATGTTCGAAACAGAGCATCATAC |
| DWF4_R | GGGGACCACTTTGTACAAGAAAGCTGGGTTCAGAATACGAGAAACCCTAATAG |
| CPD_F | GGGGACAAGTTTGTACAAAAAAGCAGGCTTTATGGCCTTCACCGCTTTTC |
| CPD_R | GGGGACCACTTTGTACAAGAAAGCTGGGTTAGTAGCAAAATCACGGCGCT |
| ROT3_1_F | GGGGACAAGTTTGTACAAAAAAGCAGGCTTTATGCAACCTCCGGCAAGC |
| ROT3_1_R | GAGAATAGAGTTTTCTCCAA |
| ROT3_2_F | GTCCCTGCATACCCTAAATC |
| ROT3_2_R | CAATTCCAATTTACGCCTCT |
| ROT3_3_F | TTTATTTATCTACAGGAGGAG |
| ROT3_3_R | GGGGACCACTTTGTACAAGAAAGCTGGGTTATGATCTTCAAGTGAGATCGGAG |
| CYP90D1_F | GGGGACAAGTTTGTACAAAAAAGCAGGCTTTATGGACACTTCTTCTTCAC |
| CYP90D1_R | GGGGACCACTTTGTACAAGAAAGCTGGGTTTATTCTTTTGATCCAAATGG |
| BR6OX1_F | GGGGACAAGTTTGTACAAAAAAGCAGGCTTTATGGGAGCAATGATGGTG |
| BR6OX1_R | GGGGACCACTTTGTACAAGAAAGCTGGGTTGTAGGGTGAAATCCTAAGATGG |

| | |
|---|---|
| BR6OX2_F | GGGGACAAGTTTGTACAAAAAAGCAGGCTTTATGGGCATAATGATGATGATTTTGGG |
| BR6OX2_R | GGGGACCACTTTGTACAAGAAAGCTGGGTTGTAAGGTGAACACTTAAGATGGTATCC |
| BR6OX2_noSP_F | GGGGACAGCTTTCTTGTACAAAGTGGGGATGCGATATTCTAAGAAAGGTC |
| BR6OX2_R_ stop | GGGGACAACTTTGTATAATAAAGTTGATCAGTAAGGTGAACACTTAAGATG |
| SP-GFP_F | GGGGACAAGTTTGTACAAAAAAGCAGGCTTTATGGGCATAATGATGATGATTTTGGGTCTTCTTGTGATCATTGTTTGTTTATGTACTGCTCTTCTCCGATGGAACCAGATGGTGAGCAAGGGCGAG |
| SP-GFP_R | RV: GGGGACCACTTTGTACAAGAAAGCTGGGTTCTTGTACAGCTCGTCCATGC |
| BAS1_F | GGGGACAAGTTTGTACAAAAAAGCAGGCTTTATGGAGGAAGAAAGTAGCAGCTGG |
| BAS1_R | GGGGACCACTTTGTACAAGAAAGCTGGGTTATCCTCATGATTGGTCAATCTCCG |
| **RT-qPCR primers** | |
| DWF4_Fw | GTGATCTCAGCCGTACATTTGGA |
| DWF4_Rv | CACGTCGAAAAACTACCACTTCCT |
| CPD_Fw | CCCAAACCACTTCAAAGATGCT |
| CPD_Rv | GGGCCTGTCGTTACCGAGTT |
| ROT3_Fw | GGAAGCTTAGGCTGGCCG |
| ROT3_Rv | CCCGTATAAAGACTTTCGTTTGTCC |
| BR6OX1_Fw | CGTGACTTGGAAATCAACGGT |
| BR6OX1_Rv | ACTCCAAGCTCTTCTTCATCCA |
| BR6OX2_Fw | ACCCATGGAGATGGATGGAAAA |
| BR6OX2_Rv | CAGGGCAAAGCCTAACTCCA |
| BAS1_Fw | TTGGCTTCATACCGTTTGGC |
| BAS1_Rv | TTACAGCGAGTGTCAATTTGGC |
| ACTIN2_Fw | GATGAGGCAGGTCCAGGAATC |
| ACTIN2_Rv | AACCCCAGCTTTTTAAGCCTTT |
| **Genotyping primers** | |
| DWF4_L | GGAAACAAAGAACAGACGATGATC |
| DWF4_R | GTTGCCATCTCCAAGGATTAAAG |
| CPD_L | AAGGTCCTACTTTATGCAGAA |
| CPD_R | AAAAACATAAGTGAGAAGGCCGAAT |
| CPD_ins | GCACGAGGGAGCTTCCA |
| BR6OX1_L | CAAGAAATAACATTGATAGGTTGCTAAG |
| BR6OX1_R | CATGGATTAAAGATCAATGGGTC |
| BR6OX2_L | GACTTTGAAAAAACCAGAGGTTG |
| BR6OX2_R | GAGTTCTAAGTCGTGAGTAGTTTTC |
| LB1.3 for SALK mutants | ATTTTGCCGATTTCGGAAC |
| ROT3_g_Fw | ATGCAACCTCCGGCAAG |
| ROT3_g_Rv | GTTTGTTATAACCCCTAATACTAATG |
| ROT3_C_Fw | GTCCCTGCATACCCTAAATC |
| ROT_C_Rv | CCTCCACAACCTTTTTTACCATC |
| CYP90D1_L | ATGGACACTTCTTCTTCAC |
| CYP90D1_R | TATTCTTTTGATCCAAATGG |

| | |
|---|---|
| FISH2 for *cyp90d* mutant | CAGTCATAGCCGAATAGCCTCTCCA |

**Supplementary Video 1.** Long-term confocal imaging of 6-day-old *pDWF4WF4-GFP/dwf4* roots by vertical-stage microscopy.

**Supplementary Video 2.** Long-term confocal imaging of 6-day-old *pROT3:ROT3-GFP/rot3* roots by vertical-stage microscopy.

**Supplementary Video 3.** Long-term confocal imaging of 6-day-old *pCPD:CPD-GFP/cpd* roots by vertical-stage microscopy.

**Supplementary Video 4.** Long-term confocal imaging of 6-day-old *pBES1:BES1-GFP* roots by vertical-stage microscopy.
